# Mechanism by which PF-3758309, a Pan Isoform Inhibitor of p21-Activated Kinases, Blocks Reactivation of HIV-1 Latency

**DOI:** 10.1101/2022.08.02.502581

**Authors:** Benni Vargas, James Boslett, Nathan A. Yates, Nicolas Sluis-Cremer

## Abstract

The “block and lock” strategy is one approach that might elicit a sterilizing cure for HIV-1 infection. The “block” refers to a compound’s ability to inhibit latent HIV-1 proviral transcription, while the “lock” refers to its capacity to induce permanent proviral silencing. We identified PF-3758309, a pan-isoform inhibitor of p21-activated kinases (PAKs), as a potent inhibitor of HIV-1 latency reversal (Vargas *et al*., Antimicrob Agents Chemother. 2019;63(2):e01744-18). The goal of this study was to define the mechanism(s) involved. We found that both 24ST1NLESG cells (a cell line model of HIV-1 latency) and purified CD4+ naïve and central memory T cells express high levels of PAK2, and lower levels of PAK1 and PAK4. Knockdown of PAK1 or PAK2, but not PAK4, in 24ST1NLESG cells resulted in a modest, but statistically significant decrease in the magnitude of HIV-1 latency reversal. Overexpression of PAK1 significantly increased the magnitude of latency reversal. A phospho-protein array analysis revealed that PF-3758309 down-regulates the NF-κB signaling pathway, which provides the most likely mechanism by which PF-3758309 inhibits latency reversal. Finally, we used cellular thermal shift assays combined with liquid chromatography and mass spectrometry to ascertain whether PF-3758309 off-target binding contributed to its activity. In 24ST1NLESG cells and in peripheral blood mononuclear cells, PF-3758309 bound to mitogen-activated protein kinase 1 and protein kinase A; however, knockdown of either of these kinases did not impact HIV-1 latency reversal. Collectively, our study suggests that PAK1 and PAK2 play a key role in the maintenance of HIV-1 latency.

**IMPORTANCE:** The persistence of latent, replication-competent HIV-1 proviruses in resting CD4+ T cells, and other cellular reservoirs, represents a major barrier to a cure. The “block and lock” strategy is one approach proposed to elicit a sterilizing cure for HIV-1 infection. In this study, we define the mechanism by which PF-3758309, a pan-isoform p21-activated kinase (PAK) inhibitor, blocks the reversal of HIV-1 latency. Our data show that PAK1 and PAK2 play a role in the maintenance of HIV-1 latency, and further suggest that PAK inhibitors, such as PF-3758309, could form part of “block and lock” therapeutic strategies.

## INTRODUCTION

Despite effective antiretroviral therapy (ART), a reservoir of latent, replication-competent proviral HIV-1 DNA persists in resting CD4+ T cells that represents a major barrier to either a sterilizing or functional cure for HIV-1 infection (1,2). While a sterilizing cure involves the complete elimination of the latent replication-competent proviral HIV-1 DNA in the body, a functional cure results in the long-term control of HIV-1 replication in the absence of ART. The “block and lock” therapeutic strategy represents one approach for a functional cure, and involves durable epigenetic silencing of latent, replication-competent proviral HIV-1 DNA in infected CD4+ T cells by either small molecules or oligonucleotides (3,4).

Signaling pathways play a key role in HIV-1 latency (5). In a recent study, our laboratory used the 24ST1NLESG cell line model of HIV-1 latency to screen a library of structurally diverse, medicinally active, cell permeable kinase inhibitors, which target a wide range of signaling pathways, to identify inhibitors of HIV-1 latency reversal (6). The screen was carried out in the absence or presence of three mechanistically distinct latency-reversing agents, namely, prostratin, panobinostat, and JQ-1. We identified 12 kinase inhibitors that blocked the reversal of HIV-1 latency, irrespective of the latency reversal agent used in the screen. Of these, PF-3758309 was found to be the most potent. The 50% inhibitory concentrations in the 24ST1NLESG cells ranged from 0.1 to 1 nM (selectivity indices, >3,300). PF-3758309 was also found to inhibit latency reversal in CD4+ T cells isolated from HIV-1-infected donors on ART. The mechanism by which PF-3758309 inhibited reactivation of latent HIV-1 was not elucidated in this study.

PF-3758309 (**Fig. 1**) is a potent, ATP-competitive, pyrrolopyrazole inhibitor of the p21-activated kinases (PAKs) with 50% inhibitory concentrations (IC_50_) of 13.7, 190, 99, 18.7, 18.1 and 17.1 nM against PAK 1, 2, 3, 4, 5 and 6 in cell-free assays, respectively (7). PAKs are a family of serine/threonine-specific protein kinases that act as downstream effectors of the small GTPases Cdc42 and Rac, mediating an important subset of the signaling activities of these enzymes, and aberrant PAK signaling has long been associated with a number of human diseases (8). Furthermore, there is emerging evidence that PAKs also play a major role in the entry, replication and spread of many important pathogenic human viruses, including HIV-1 (9–12). In regard to HIV-1, the viral accessory protein Nef specifically associates with PAK2, and this interaction appears to be critical for several aspects of HIV-1 biology, not only directly affecting virus replication but also indirectly promoting virus spread and persistence via disturbance of key players in the antiviral immune response (10,12). The primary objective of this study was to determine how inhibition of the different PAK isoforms by PF-3758309 modulates HIV-1 latency.

**Figure 1.**
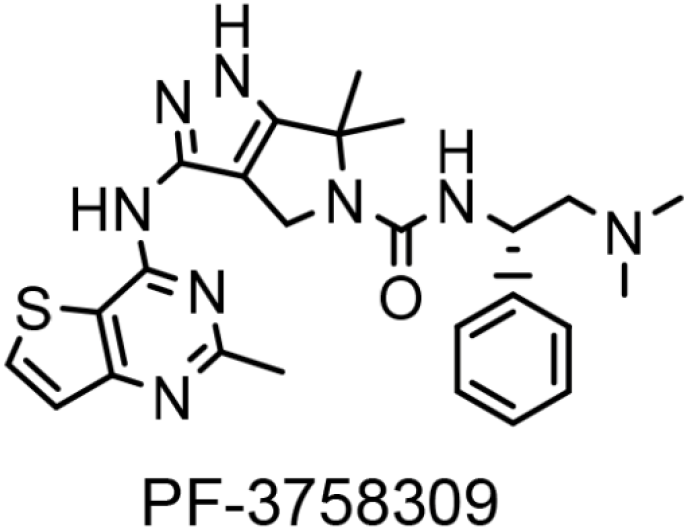
Chemical structure of PF-3758309.

## RESULTS

### Inhibition of HIV-1 latency reversal by PF-3758309 and other PAK inhibitors

PF-3758309 is an orally available, potent, ATP-competitive, pyrrolopyrazole inhibitor of the PAKs, which are involved in physiological processes including motility, survival, mitosis, transcription and translation (8). Depending on structural and functional similarities, the six members of the PAK family are divided into two groups with three members in each group (8). Group I PAKs (1–3) are activated by extracellular signals through GTPase-dependent and GTPase-independent mechanisms. In contrast, group II PAKs (4–6) are constitutively active. PF-3758309 exhibits inhibitory activity toward all of the PAK isoforms. In an attempt to delineate which PAKs contribute to the maintenance of HIV-1 latency, we first assessed the inhibitory activity of a panel of different PAK inhibitors with different isoform specificities in 24ST1NLESG cells using four mechanistically distinct latency reversal agents, namely prostratin (a protein kinase C inhibitor), panobinostat (a histone deacetylase inhibitor), JQ-1 (a bromodomain-containing protein 4 inhibitor) and TNFα (**Table 1**). In the 24ST1NLESG cell line, the integrated HIV-1 genome encodes a secretable alkaline phosphatase (SEAP) gene in the viral *env* gene, which serves as an indicator of late gene expression and was used as a measure of latency reversal in this study (13). In general, none of these PAK inhibitors demonstrated robust inhibition of HIV-1 latency reversal at concentrations that did not result in cellular cytotoxicity.

**Table 1:** Inhibition of HIV-1 latency reversal in 24ST1NLESG cells by PAK inhibitors. Data include 50% inhibitory concentration (IC_50_) values for inhibition of HIV-1 latency reversal mediated by prostratin, panobinostat, JQ-1 and TNF-α; 50% cytotoxicity concentrations (CC_50_); and selectivity indices (CC_50_/IC_50_). Data are shown as the mean ± standard deviation from at least 3 independent biological replicates.

| Inhibitor | PAK specificity | Cytotoxicity<br>CC<sub>50</sub> (μM) | IC<sub>50</sub> for Inhibition of latency reversal (μM)<br>(Selectivity Index (CC<sub>50</sub>/IC<sub>50</sub>)) | | | |
| --- | --- | --- | --- | --- | --- | --- |
| | | | Prostratin | Panobinostat | JQ-1 | TNF-α |
| PF-3758309 | PAK1/2/3/4/5/6 | 4.3 ± 1.2 | 0.00007±0.00004<br>(64529.1) | 0.0004±0.0001<br>(10070.4) | 0.0010±0.0003<br>(3364.7) | 0.00082±0.00009<br>(5265.5) |
| IPA3 | PAK1 | > 16.7 | > 16.7 | > 16.7 | 16.0 | 15.2 |
| FRAX597 | PAK1/2/3 | 1.9 | > 2.0 | >2.0 | >2.0 | >2.0 |
| FRAX486 | PAK1/2/3/4 | 0.8 | 0.1<br>(8) | >0.8 | >0.8 | > 0.8 |
| KPT9274 | PAK4 | 0.3 | > 0.3 | > 0.3 | > 0.3 | > 0.3 |
| GNE2861 | PAK4/5/6 | 49.2 | > 50 | > 50 | > 50 | > 50 |

### PAK isoform expression in 24ST1NLESG cells and in purified CD4+ naïve (T_N_) and central memory (T_CM_) T cells

PAKs are widely expressed in a variety of tissues and cells, and are often overexpressed in multiple cancer types. We used quantitative PCR or population RNA sequencing data to assess PAK isoform expression in 24ST1NLESG cells and purified CD4+ T_N_ and T_CM_ cells (**Fig. 2**). Of note, CD4+ T_N_ and T_CM_ cells are regarded as major reservoirs of latent HIV-1 infection in infected individuals on ART (1,2). In the 24ST1NLESG cell line, we noted robust expression of PAK2, and somewhat lower expression levels of PAK1 and PAK4 (**Fig. 2a**). Expression of PAK3 and PAK6 was exceptionally low, and we did not detect expression of PAK5 (**Fig. 2a**). The PAK isoform expression levels in CD4+ T_N_ and T_CM_ cells were similar to that of the 24ST1NLESG cell line (**Fig. 2b**).

**Figure 2.**
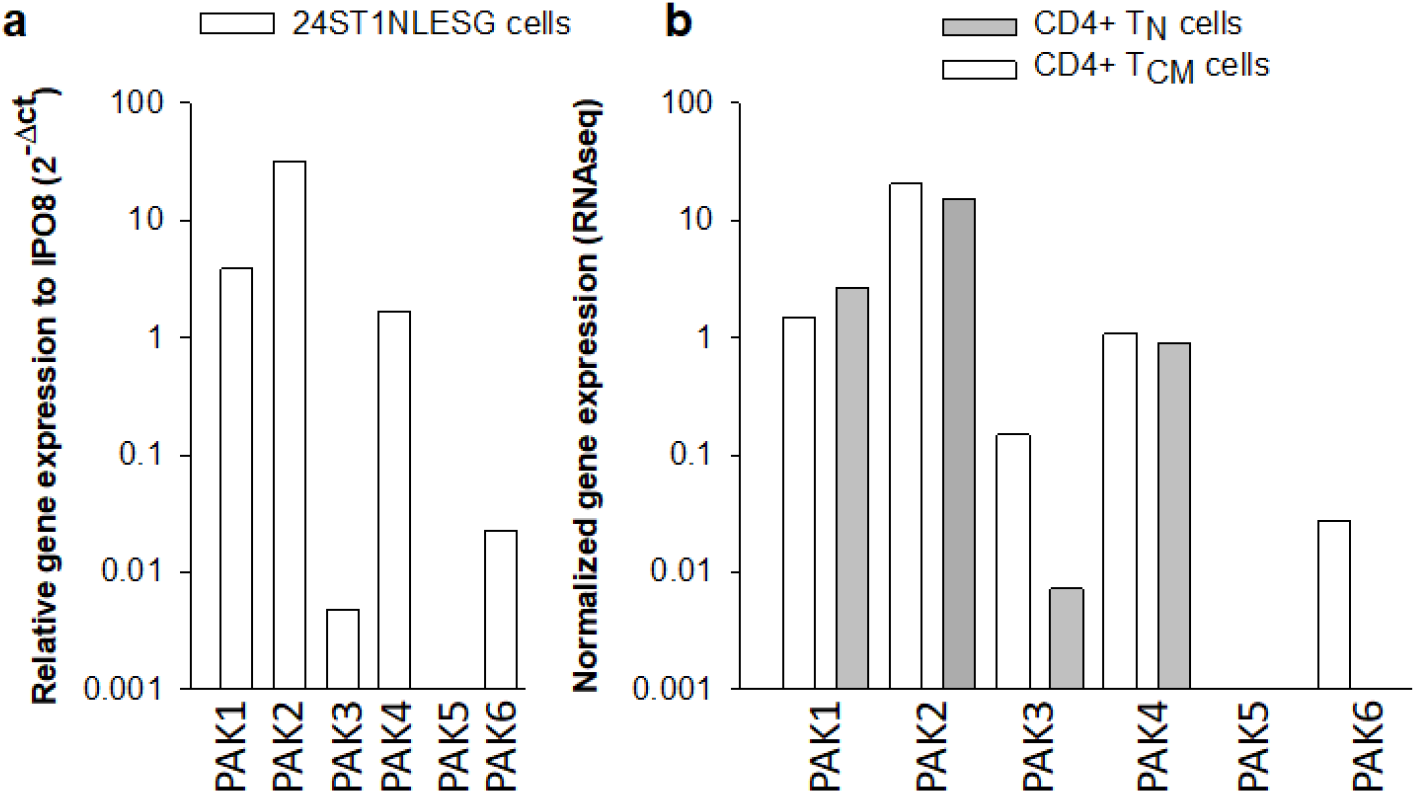
RNA expression levels of different PAK isoforms in 24ST1NLESG cells (a) and in purified naïve (T_N_) and central memory (T_CM_) CD4+ T cells. PAK RNA expression is reported relative to cellular IPO8 expression.

### Knockdown of PAK1, PAK2 and PAK4 in 24ST1NLESG cells

We used siRNA to knock down expression of PAK1, PAK2 and PAK4 in 24ST1NLESG cells (**Fig. 3**). A scrambled siRNA sequence was used as a negative control in these experiments. The magnitude of the siRNA-mediated gene silencing was confirmed 120 h post-siRNA transfection by Western blot analysis (**Fig. 3a, b**). Knockdown of PAK1, PAK2 or PAK4 did not result in any measurable cellular cytotoxicity (data not shown). Next, we assessed the effect of PAK isoform knockdown on the reversal of HIV-1 latency in 24ST1NLESG cells (72 h post-siRNA transfection) by addition of 50 ng/mL tumor necrosis factor α (TNFα) for 48 h (**Fig. 3c**). Knockdown of PAK1 and PAK2 resulted in a modest, but significant, decrease in SEAP expression. Knockdown of PAK4 did not impact reactivation of latent HIV-1 expression. These data provide evidence that inhibition of PAK1 and PAK2 contribute to the maintenance of latent HIV-1 infection in 24ST1NLESG cells.

**Figure 3.**
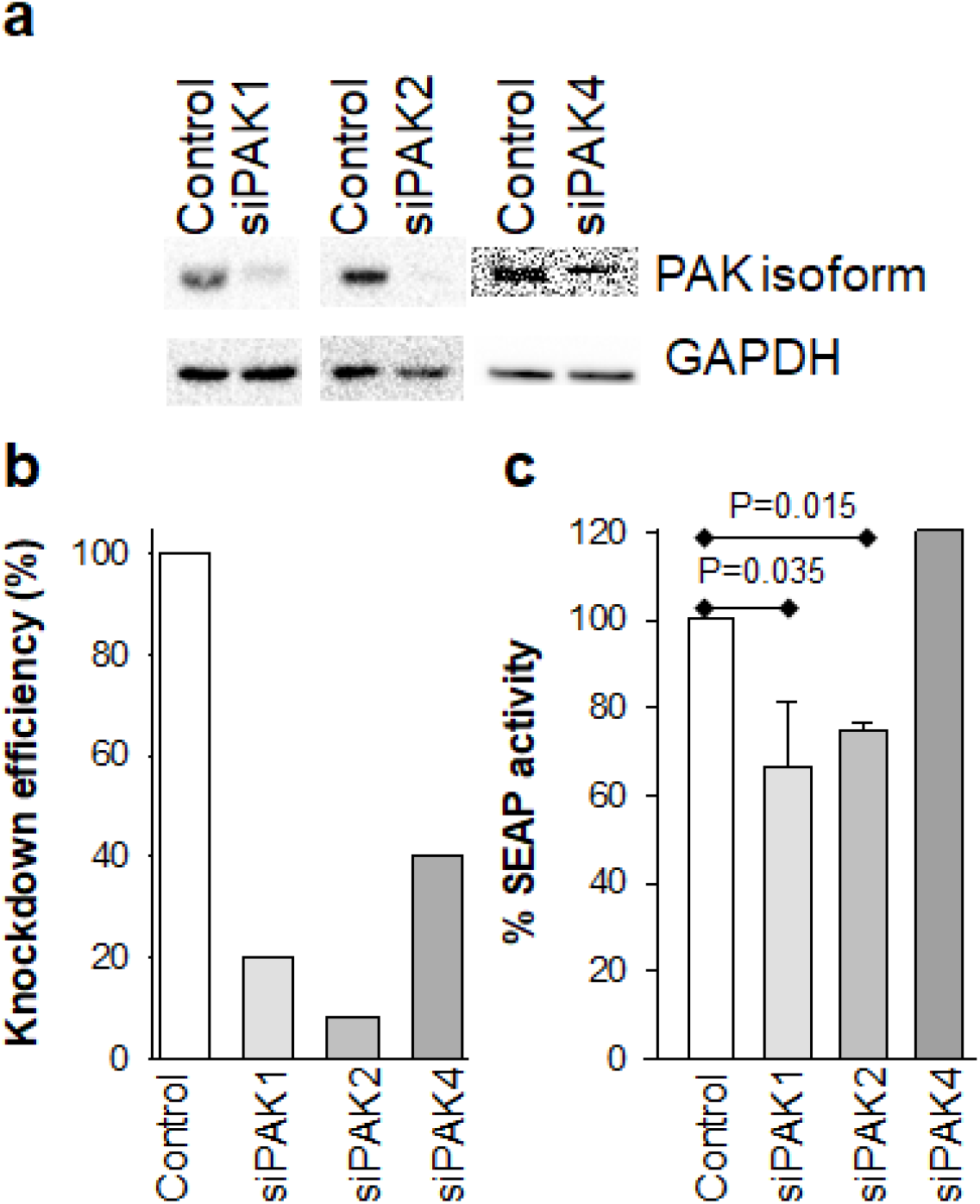
Effect of PAK1, PAK2 and PAK4 knockdown of HIV-1 latency reversal in 24ST1NLESG cells. (**a**) Western blot analysis of PAK knockdown; (**b**) Quantification of PAK knockdown using ImageJ; (**c**) effect of PAK knockdown on HIV-1 latency reversal mediated by TNF-α in 24ST1NLESG cells. Data are shown as the mean ± standard deviation from at least 3 independent biological replicates. Statistical significance evaluated using Mann-Whitney U test.

### Overexpression of PAK1 and PAK2 in 24ST1NLESG cells

To further delineate the roles of PAK1 and PAK2 in the maintenance of HIV-1 latency, we overexpressed these isoforms in 24ST1NLESG cells (**Fig. 4**). Cells were harvested 120 h post-transfection to assess overexpression by Western blot analysis. Compared to the control experiment which included cells transformed with an empty vector, we observed notable over-expression of PAK1 in the in 24ST1NLESG cells (**Fig. 4a, b**). By contrast, we could not assess PAK2 overexpression due to the experimental difficulty in purifying high plasmid yields in bacteria, as reported previously (14,15). Overexpression of PAK1 in 24ST1NLESG cells resulted in a statistically significant increase in SEAP expression (**Fig. 4c**), providing additional evidence that this isoform contributes to the maintenance of HIV-1 latency.

**Figure 4.**
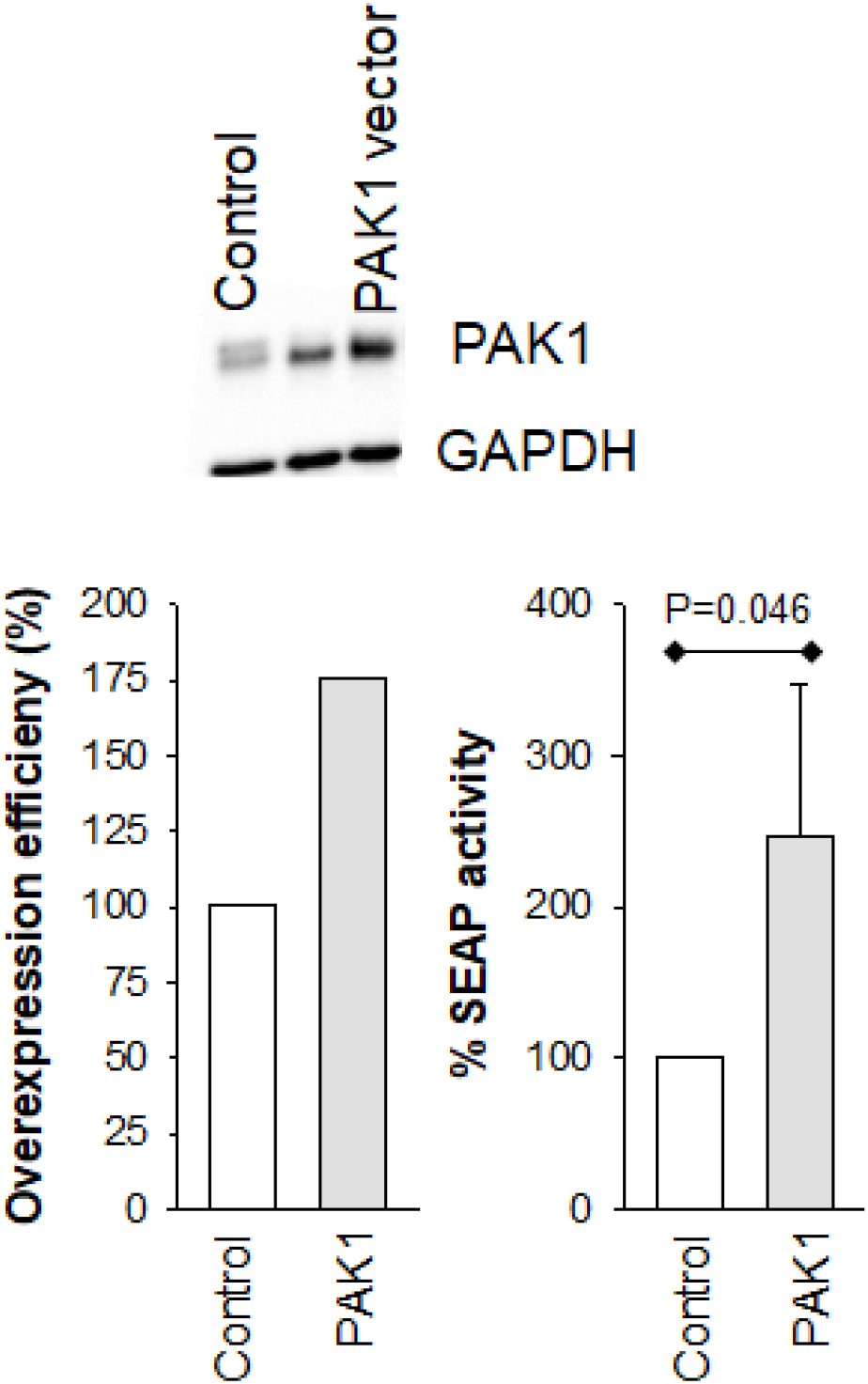
Effect of PAK1 overexpression on HIV-1 latency reversal in 24ST1NLESG cells. (**a**) Western blot analysis of PAK1 overexpression; (**b**) Quantification of PAK1 overexpression using ImageJ; (**c**) effect of PAK1 overexpression on HIV-1 latency reversal mediated by TNF-α in 24ST1NLESG cells. Data are shown as the mean ± standard deviation from at least 3 independent biological replicates. Statistical significance evaluated using Mann-Whitney U test.

### PAK signaling and HIV-1 latency reversal

PAKs act as key signal transducers in several signaling pathways, including Ras, Raf, NFκB, Akt, Bad and p53. Protein phosphorylation plays an important role in cell signaling. Accordingly, we used the Phospho Explorer Antibody Array (Full Moon Biosystems) to quantitate broad-scope phosphorylation profiling changes in 24ST1NLESG cells exposed to 8 nM PF-3758309 or DMSO (control) for 24 h to try to better understand how inhibition of PAKs by PF-3758309 controls HIV-1 latency. The Phospho Explorer Antibody Array conducts phosphorylation profiling with 1318 site-specific antibodies from over 30 signaling pathways. In our data analysis plan, we only considered phosphorylation changes that exhibited fold changes in phosphorylation that were > 1.4 and < 0.6 between control and PF-3758309 treated cells. In total, we identified 47 proteins that exhibited site-specific increased phosphorylation and 14 proteins that exhibited site-specific decreased phosphorylation in PF-3758309 treated cells, respectively (**Supplementary Table 1**). Next, we used the Ingenuity^®^ Pathway Analysis to identify which signaling pathways were regulated by PF-3758309 (**Fig. 5**). This analysis revealed down-regulation of the NF-κB pathway (p-value 2.8 × 10^−24^), which is consistent with prior studies demonstrating that PF-3758309 suppressed NF-κB signaling in lung cancer cells (16,17). Importantly, multiple studies have demonstrated that both the canonical and non-canonical NF-κB signaling pathways drive HIV-1 proviral expression (18). As such, down-regulation of the NF-κB pathway by PF-3758309 provides the most likely mechanism by which the drug inhibits latency reversal.

**Figure 5.**
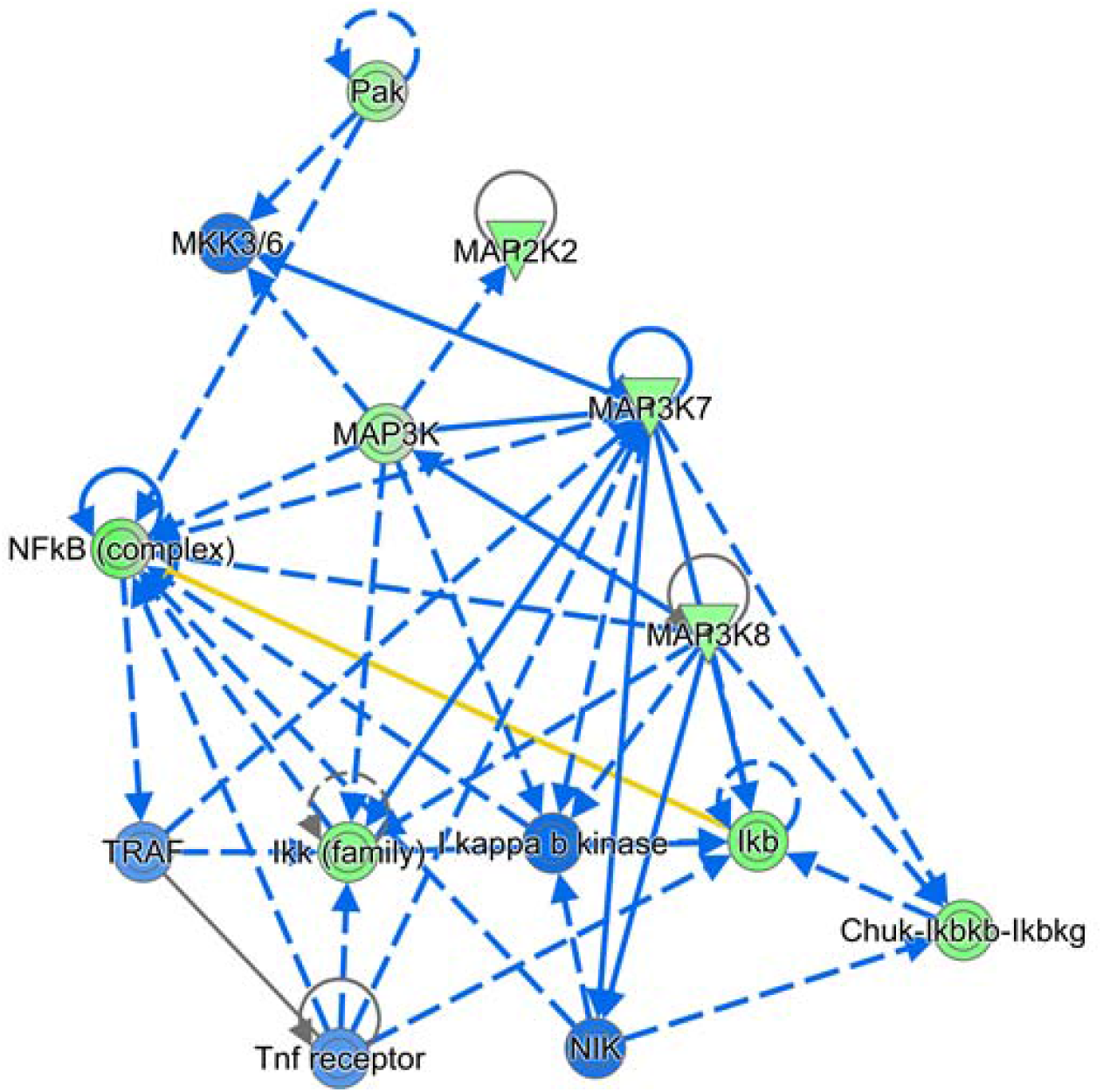
Network of signaling pathways impacted in 24ST1NLESG cells by PF-3758309 treatment. Qiagen Ingenuity Pathway Analysis was used to generate the functional network. NF-κB signaling pathway was found to be significantly down-regulated (p-value 2.83 × 10^−24^). Lines and arrows between nodes represent direct (solid lines) and indirect (dashed lines) interactions between molecules as supported by information in the Ingenuity knowledge base. Green nodes represent decreased measurement in the dataset. Blue color represents predicted inhibition. Yellow line indicates findings inconsistent with state of downstream molecule. Gray line indicates effect not predicted. Circular line arrow indicates molecule interacts with itself.

### Potential off-target effects of PF-3758309

Kinase inhibitors, such as PF-3758309, can produce off-target effects by inducing changes in molecules other than the one specifically targeted. Such off-target effects are generally attributed to non-specific binding or to cross-talk. PF-3758309 is an exceptionally potent inhibitor of HIV-1 latency reversal, with an IC_50_ for inhibition of latency reversal ranging from 0.070-1 nM depending on the latency reversing agent used (**Table 1**). As described above, knockdown or overexpression of PAK1 or PAK2 only provided a modest effect on latency reversal, possibly suggesting that the drug may bind to and inhibit other proteins also involved in the maintenance of HIV-1 latency. To identify possible cellular off-targets for PF-3758309 we used cellular thermal shift assays (CETSA) combined with liquid chromatography and mass spectrometry (LC-MS/MS). CETSA relies on the principle that small molecule binding to a protein alters the melting behavior of that protein, typically increasing it (19). Briefly, CETSA involves treating cellular protein extracts with a test drug or its vehicle followed by heating to induce protein denaturation. Protein aggregates are pelleted by centrifugation, and the protein in the supernatant is proteolytically digested to peptides which are quantified by LC-MS/MS analysis. Peptides enriched in the drug-treated extracts represent potential binding partners for the drug of interest, as these were stabilized by drug treatment. We prepared 1 mg/ml cytosolic extracts of 24ST1NLESG cells or peripheral blood mononuclear cells (PBMC), added vehicle (i.e., DMSO) or 10 μM PF-3758309 and performed CETSA at 3 temperatures (50, 55, and 60°C). LC-MS/MS was performed on 6 replicates from each of the conditions noted above. In some experiments, a decoy vehicle group was analyzed (vehicle vs. vehicle) to determine the statistical significance threshold for real “hits” in the experimental group (PF-3758309 vs. vehicle). **Fig. 6** and **Supplementary Fig. 1** show series of volcano plots for the 55°C data in 24ST1NLESG cells (**Fig. 6**) and peripheral blood mononuclear cells (**Supplementary Fig. 1**), respectively. The vehicle vs. vehicle decoy comparison (panels 1a and 2a) show similar thresholds required for statistical significance (~P<0.001). From this, we established a practical filter for determining robust thermal stabilization events with PF-3758309 in the drug vs. vehicle comparison (panels **1b** and **2b**). This practical filter specified that, in order to be considered a robust thermal stabilization, the protein must have at least two peptides with levels statistically different (P<0.001) between the PF-3758309 and vehicle-treated samples, as there were no proteins meeting this requirement in the vehicle vs. vehicle decoy group. Using this filter, we found two off-target binding proteins of PF-3758309, mitogen-activated protein kinase 1 (MAPK1) and protein kinase A (PKA). Interestingly, this finding was common to both cell types, as well as the analysis at 60°C (not shown). Thus, the finding repeated across several independent experiments with varying proteomes and at different protein denaturation temperatures. Next, we used siRNA to knock down expression of MAPK1 and PKA in 24ST1NLESG cells (**Fig. 7**). The magnitude of the siRNA-mediated gene silencing was confirmed 120 h post-siRNA transfection by Western blot analysis (**Fig. 7a, b**). However, knockdown of MAPK1 and PKA did not impact HIV-1 latency reversal in 24ST1NLESG cells (**Fig. 7c**), suggesting that off-target binding of PF-3758309 to these proteins does not contribute to the activity of the drug.

**Figure 6:**
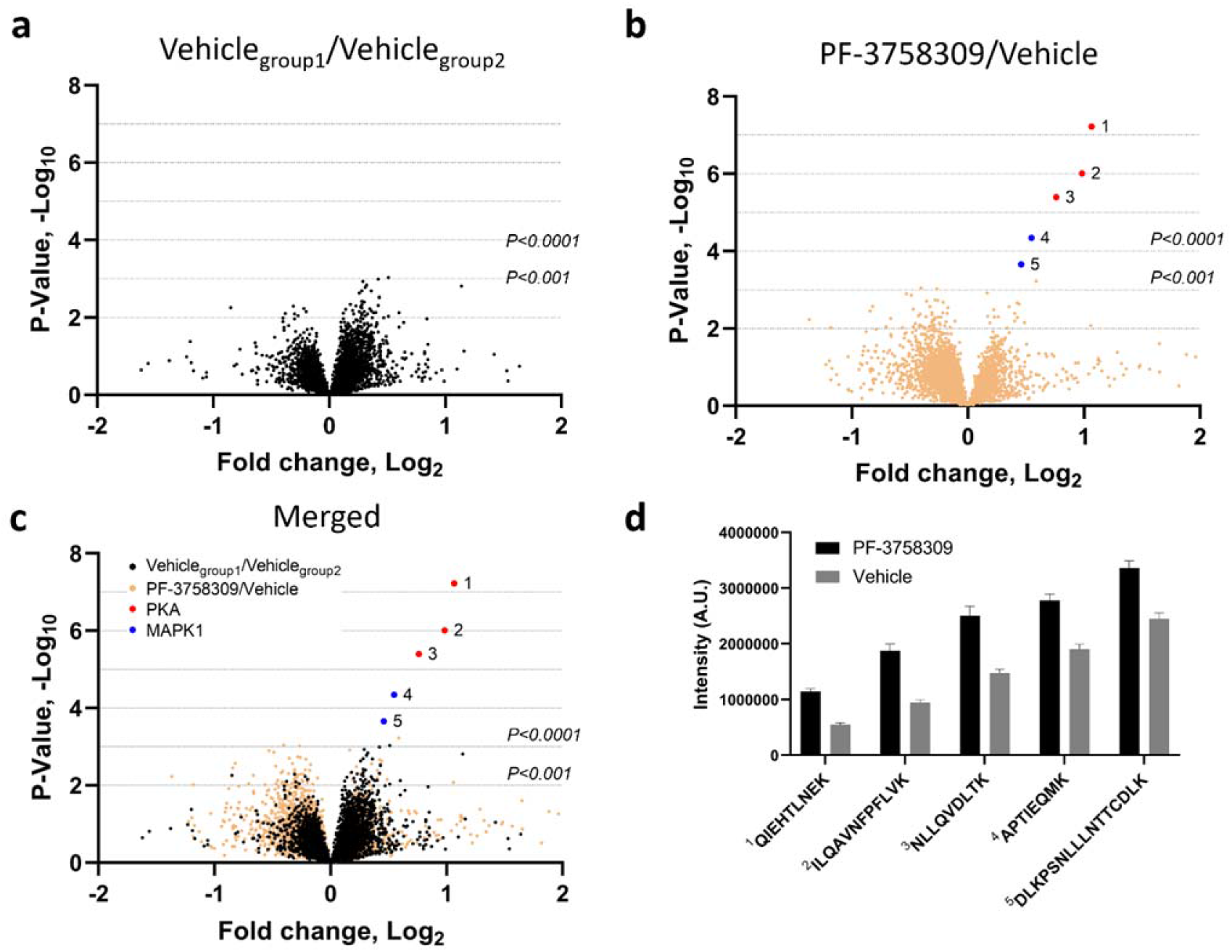
Drug/protein binding analysis of PF-3758309-treated 24ST1NLESG cells. Volcano plots for (a) two identical vehicle groups (decoy comparison), (b) PF-3758309 and vehicle groups (experimental comparison), and (c) the decoy and experimental comparisons merged. The x-axis indicates the relative fold change (log_2_-transformed expression values of peptides in the absence or presence of PF-3758309) and the y-axis indicates the -log10 of the p values. (**d**) Bar graph illustrating enrichment of PKA and MAPK1 peptides in 24ST1NLESG cell lysates treated with PF-3758309. Bar graph data are shown as the mean ± standard error.

**Figure 7.**
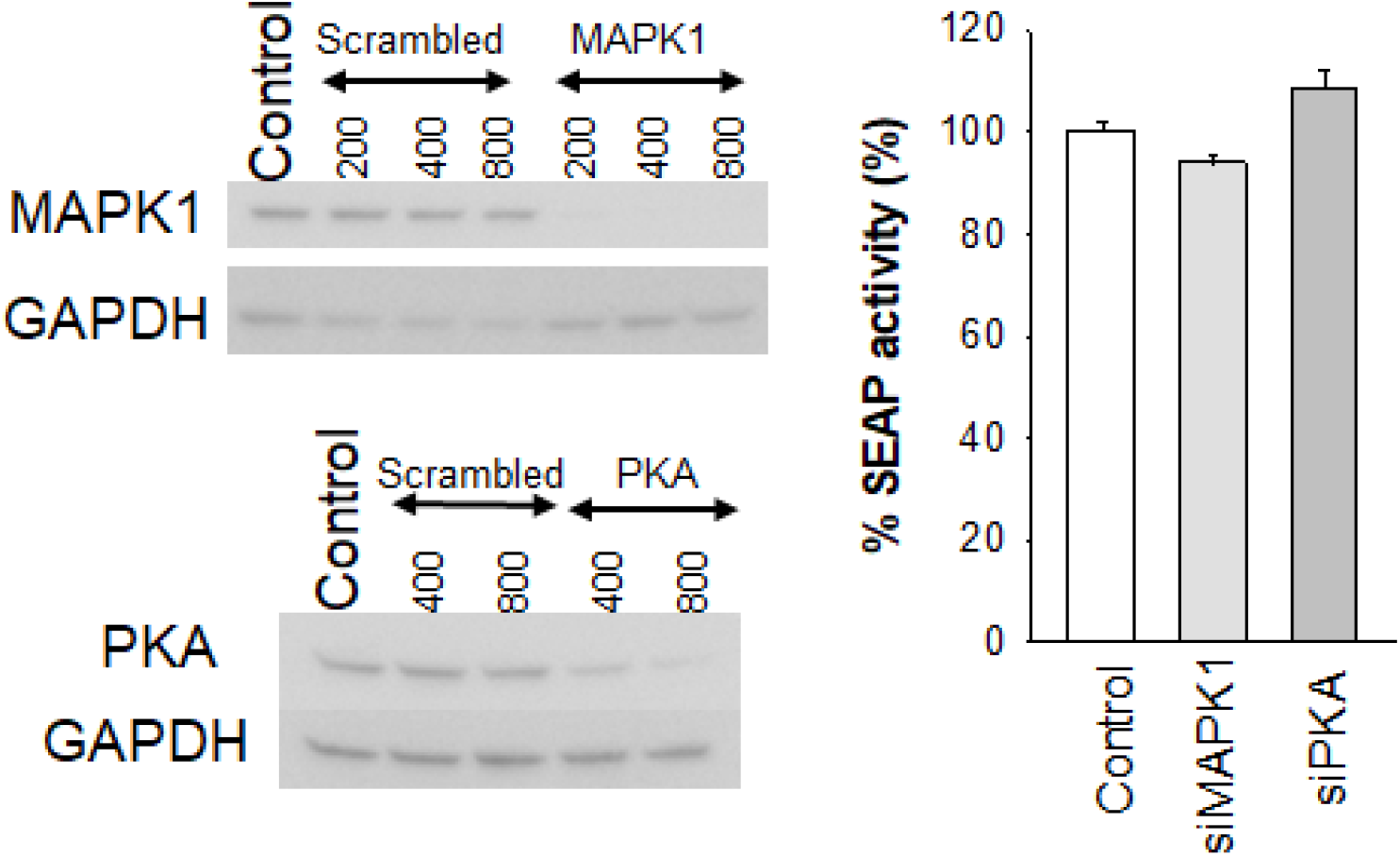
Effect of MAPK1 and PKA knockdown of HIV-1 latency reversal in 24ST1NLESG cells. (**a**) Western blot analysis of MAPK1 knockdown using different siRNA concentrations; (**b**) Western blot analysis of PKA knockdown using different siRNA concentrations; (**c**) effect of MAPK1 and PKA knockdown on HIV-1 latency reversal mediated by TNF-α in 24ST1NLESG cells. Data are shown as the mean ± standard deviation from at least 3 independent biological replicates. Statistical significance evaluated using Mann-Whitney U test.

## DISCUSSION

The persistence of latent, replication-competent HIV-1 proviruses in resting CD4+ T cells represents a major barrier to curing HIV-1 infection (1,2). To date, efforts to eradicate this viral reservoir via the shock and kill approach have not led to complete, long term viral suppression in either cell or animal models. Thus, we need to consider alternate approaches that could lead to a sterilizing or functional cure for HIV-1 infection. The block and lock approach seeks to silence the transcriptional activity of latent proviruses, such that when ART is removed viral rebound is significantly delayed or, better yet, prevented (3,4). Several research groups have identified small molecule inhibitors that target different factors of the HIV-1 transcription machinery, leading to a block and lock phenotype (20). However, blocking only one transcription pathway may not be sufficient to silence all proviruses, and thus it is likely that successful implementation of this strategy will require a combination of inhibitors. In this regard, there is a critical need to identify new molecules with different mechanisms of action. Our laboratory recently discovered that the pan isoform PAK inhibitor PF-3758309 is an exceptionally potent inhibitor of HIV-1 latency reactivation (IC_50_ in the pM to low nM range) with a huge selectivity index (> 3,000) (6). In this study, we sought to define the role of different PAK isoforms in the maintenance and reversal of HIV-1 latency, and to elucidate how PF-3758309 blocks HIV-1 latency reversal.

Our data show that CD4+ T cells express high levels of PAK2, and lower levels of PAK1 and PAK4. Knockdown of PAK1 or PAK2, but not PAK4, in 24ST1NLESG cells resulted in a modest, but statistically significant decrease in the magnitude of HIV-1 latency reversal. Overexpression of PAK1 significantly increased the magnitude of latency reversal. The modest changes in HIV-1 latency reversal that were observed upon knockdown or overexpression of these PAK isoforms could be due to biological (i.e., their role in the maintenance and reversal of HIV-1 latency) and/or experimental parameters. In regard to the latter, the addition of TNF-α to reverse HIV-1 latency in 24ST1NLESG cells induces T cell proliferation, and as such knockdown or overexpression of the PAK isoforms may be rapidly lost after addition of the cytokine. Interestingly, siRNA-mediated knockdown of PAK1 was previously shown to inhibit LTR-dependent transcription in human T-cell leukemia virus type 1 (HTLV-1) transformed MT4 cells and in cells transfected with an infectious clone of HTLV-1 (11). The effect size observed in that study was similar to what we report here. Furthermore, a prior study also revealed that PAK1 depletion strongly inhibited HIV-1 infection in multiple cell systems and decreased levels of integrated provirus; and that overexpression of a constitutively active PAK1 enhanced HIV-1 infection (10). Collectively, these studies highlight an important role for PAK1 and PAK2 in the maintenance of HIV-1 (and HTLV-1) latency.

PAKs act as key signal transducers in several signaling pathways, and protein phosphorylation plays an important role in cell signaling. We used the Phospho Explorer Antibody Array to quantitate broad-scope phosphorylation profiling changes in 24ST1NLESG cells exposed to PF-3758309. This analysis revealed that the inhibitor down-regulates the NF-κB signaling pathway, a finding that is consistent with prior published data (16,17). NF-κB signaling has both antagonistic and agonistic effects on HIV-1 latency reversal (21). Indeed, many HIV-1 latency reversing agents, including PKC agonists and SMAC mimetics, function by stimulating the NF-κB pathway (i.e. they act as agonists). In contrast, suppression or antagonism of the NF-κB pathway drives HIV-1 into latency. In our data set, we see decreased phosphorylation of NF-κB p65 at residue T254 (**Supplementary Table 1**). Phosphorylation of p65 at T254 stabilizes and promotes nuclear translocation of NF-κB (22). We also observed decreased phosphorylation of IKKα/β at S180/S181 and NF-κB p100/p52 at S865. Phosphorylation of IKKα/β at S180/181is associated with increased RelA nuclear translocation, acetylation, DNA binding, and transactivation activity (23), while phosphorylation of NF-κB p100/p52 at S865 is also associated with activation of the NF-κB pathway (24). Collectively, these data suggest that the primary mechanism by which PF-3758309 inhibits reversal of HIV-1 latency is via suppression of the NF-κB signaling pathway. Given that small molecule inhibitors can produce off-target effects via non-specific binding or cross-talk, we used CETSA LC-MS/MS to identify possible cellular off-targets of PF-3758309. While this analysis revealed that PF-3758309 bound to mitogen-activated protein kinase 1 and protein kinase A, knockdown of either of these kinases did not impact HIV-1 latency reversal.

Taken together, our data show that PAK1 and PAK2 play a role in the maintenance of HIV-1 latency, and further suggest that PAK inhibitors, such as PF-3758309, could form part of “block and lock” therapeutic strategies. In this regard, PF-3758309 was administered to patients with advanced solid tumors in a phase I clinical trial (ClinicalTrials.gov Identifier: NCT00932126). While the drug failed to advance due to the lack of an observed dose-response relationship, there were no safety concerns that contributed to the study termination. This suggests that PF-3758309 could re-purposed for other clinical indications, including HIV-1.

## MATERIALS and METHODS

### Reagents

The 24ST1NLESG cell line of HIV-1 latency was kindly provided by Dr. Joseph P. Dougherty (Robert Wood Johnson Medical School). Analysis of HIV-1 latency reversal in the 24ST1NLESG cells was conducted as described previously (13). All kinase inhibitors, including PF-3758309, were purchased from Selleckchem (Houston, TX). siRNA molecules were obtained from DharmaconTM (Horizon Discovery). All other reagents were of the highest quality available and were used without further purification.

### Quantification of PAK expression

RNA was extracted from purified CD4+ T_N_ and T_CM_ cells and from 24ST1NLESG cells using the RNeasy mini kit (Qiagen, Valencia, CA) according to the manufacturer’s protocol. Transcriptome analysis for the CD4+ T_N_ and T_CM_ cells was carried out by RNA sequencing (RNA-Seq; GENEWIZ, Azenta Life Sciences). The RNA from 24ST1NLESG cells was DNase-treated with RNase-Free DNase (Qiagen, Valencia, CA) and reverse transcribed using random primers with the AffinityScript multiple temperature reverse transcriptase (Agilent, Wilmington, DE). Quantitative PCR (qPCR) was performed using a Bio-Rad CFX96 real-time PCR detection system, using 2× Lightcycler 480 probes master (Roche, Indianapolis, IN) for IPO8 and Maxima SYBR Green/ROX qPCR Master Mix (2X) (Thermofisher, Waltham, MA) for PAK using primers described in **Supplementary Table 2**. Quantification for each qPCR reaction was assessed by the 2^-Δ*CT*^ algorithm, relative to IPO8 expression.

### siRNA transfection of 24ST1NLESG cells

24ST1NLESG cells were transfected by electroporation using the Neon Transfection System (Life Technologies, Carlsbad, CA). Electroporation parameters were 1350 Pulse Voltage (V), 10 Pulse Width (ms), 3 Pulse Number, 2 × 10^7^ cell/mL. Final concentrations were 1 x 10^7^ cell/mL, 1 μM siRNA (siPAK1, Dharmacon, cat # L-003521-00-0020; siPAK2, Dharmacon, cat # L-003597-00-0020; siPAK4, Dharmacon, cat # L-003615-00-0020; siMAPK1, Thermofisher cat # 4390824; siPKA, Thermofisher cat # 4390825; Non-targeting Control, Dharmacon, cat # D-001810-10-20). The efficiency of siRNA knockdown was assessed by Western blot analysis. Anti-PAK1 (cat # 2602S), anti-PAK2 (cat #2608S), anti-PAK4 (cat # 62690), anti-PKA (cat # 4782), anti-MAPK (cat # 9108), and anti-GAPDH (cat # 97166S) antibodies were purchased from Cell Signaling. The efficiency of siRNA knockdown was quantified using ImageJ.

### PAK1 and PAK2 Overexpression

Transfection of 24ST1NLESG cells was carried out with 8 μg PAK1 expression plasmid (Addgene, cat # 12208), PAK2 expression plasmid (Sino Biologicals, cat # HG10085-UT) or 8 μg empty vector (Sino Biologicals, cat # CV011). Cells were harvested 120 h post-transfection for Western blot to assess overexpression efficiency.

### Phospho Explorer Antibody Array

5×10^6^ 24ST1NLESG cells exposed to 8 nM PF-3758309 or DMSO (control) were sent to Full Moon Biosystems (Sunnyvale, CA) for analysis. In our data analysis, Phospho-ratios are changes between PF-3758309-treated vs. DMSO control where the ratio is equal to the signal intensity of the Phospho Site-Specific Antibody over the signal intensity of the site-specific antibody. Phospho-ratios of an absolute fold change > 1.4 or < 0.6 were included in the analysis. P-values were calculated using the Fisher’s Exact Test. Ingenuity®Pathway Analysis (IPA; QIAGEN, Redwood city, CA, USA) was used to further analyze the data. The software mapped each of the proteins to the repository of information in the Ingenuity Pathways Knowledge base. Molecular networks and canonical pathways regulated by PF-3758309 were obtained using IPA core analysis.

### Cellular Thermal Shift Assays

Cell lysates from 24ST1NLESGs or PBMC were treated with DMSO (vehicle) or 10 μM PF-3758309 and subjected to cellular thermal shift assays, as described previously (25). For each cell type, an individual CETSA experiment was performed at 50, 55, and 60°C to cover the majority of the mammalian proteome melting curve (26). The extracts were adjusted to 1 mg/mL protein concentrations and were divided into 50 μL aliquots (*n* = 6) of compound and vehicle-treated samples. The samples were heated in parallel at a fixed temperature for 10 min, followed by a 5-min incubation at room temperature. Samples were then centrifuged at 25,000 × *g* for 10 min at room temperature, and the supernatant denatured and digested with trypsin utilizing the FASP method (27). Samples were freeze-dried in a vacuum concentrator and resuspended in 30μl 0.1% formic acid in water. Nanoflow LC-MS/MS analysis was performed on a Waters NanoAcquity system (Milford, MA). Samples (1 μl) were injected via autosampler onto a 25 cm × 75 μM ID reversed phase column packed with 3 μM Reprosil (New Objective, Boston, MA). Peptides were separated and eluted with a gradient from 2 to 32% acetonitrile in 0.1% formic acid over 70 min at 300 nL/min into an LTQ Orbitrap XL hybrid mass spectrometer (Thermo Fisher Scientific, Waltham, MA) using a data-dependent top 8 method in positive ion mode, with spray voltage set at 2.0 kV. Full scan spectra were acquired in the range of m/z 350–1600 at 60,000 resolution using an automatic gain control target of 1 × 10^6^. Tandem mass spectra were acquired in the linear ion trap with a 35% normalized collision energy setting and an MS/MS ion target of 5 × 10^4^. Mass spectrometry data were analyzed using MaxQuant (version 1.6.7.0). Spectra were searched against the Uniprot reference database using the MaxQuant built-in peptide identification algorithm, Andromeda. Trypsin was specified as the digestion protease with the possibility of two missed cleavages. Acetylation (protein N-terminus) and oxidation of methionine were set as default variable modifications while carbamidomethylation of cysteine residues was set as a fixed modification. Other database search parameters included 20 ppm and 0.5 Da mass tolerances for precursor and product ions, respectively. Intensities for all peptides were assigned by MaxQuant using full scan mass spectra.Quantified peptides with large within replicate variability were filtered using a minimum occupancy filter (real values were required in a minimum of 3 out of 6 replicates for inclusion). The MaxQuant Peptide Groups file was then analyzed using Microsoft Excel and statistical significance was established using the Student’s *t* test on peptide peak intensity values. The GraphPad Prism software was used to generate a volcano scatterplot of the statistical significance versus magnitude of change for each protein group at a given temperature. Proteins were considered as “hits” if the relative abundances of two peptides were found with P-Values<0.001 between vehicle and PF-3758309-treated groups.

## ACKNOWLEDGMENTS

This research was supported by a grant from the National Institutes of Allergy and Infectious Diseases (R21AI57392).

## SUPPLEMENTARY DATA

**Supplementary Table 1:**
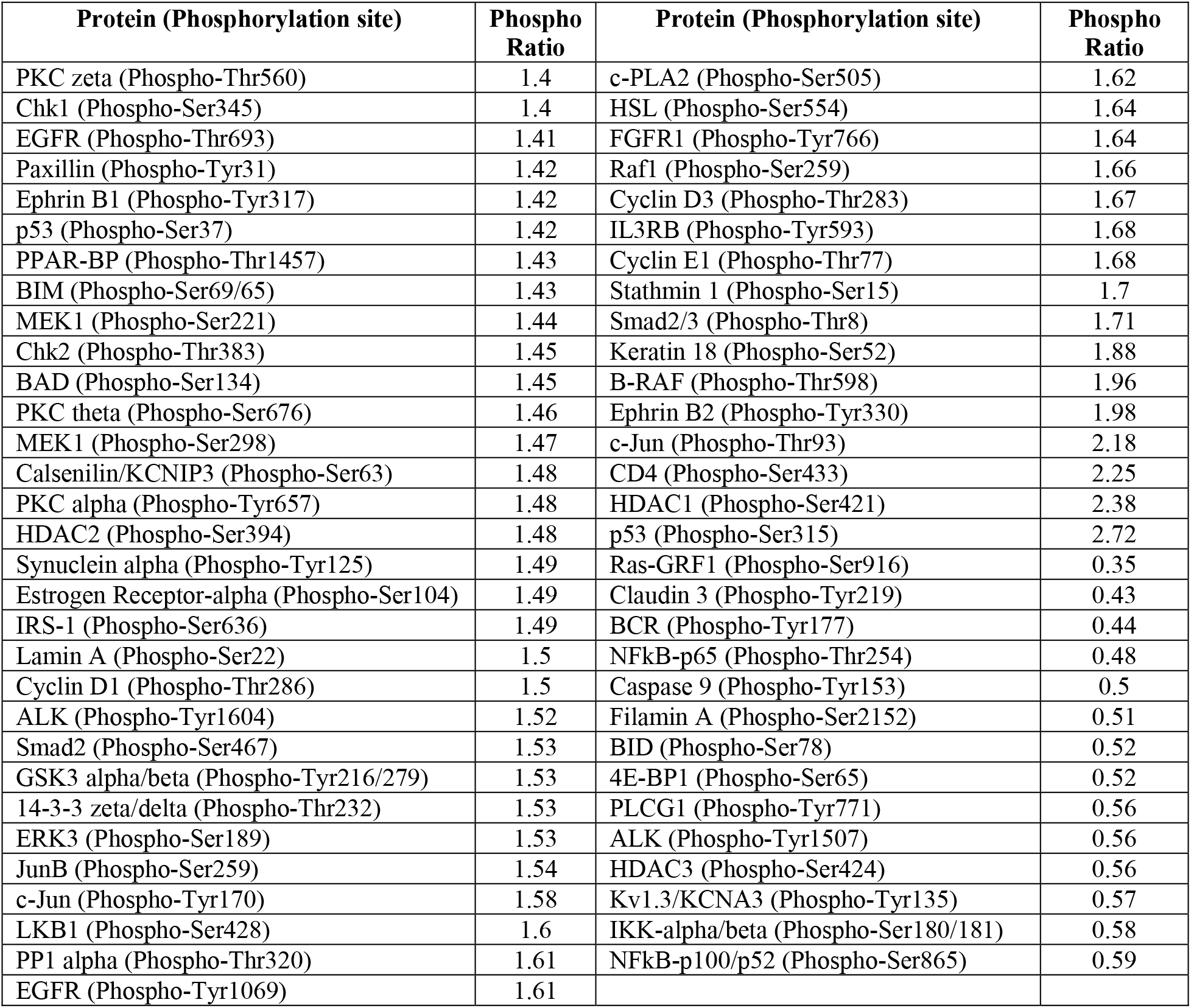
Phospho-ratios between PF-3758309-treated and DMSO-treated 24ST1NLESG cells determined from Phospho Explorer Antibody Array data. The ratio is equal to the signal intensity of the Phospho site-specific antibody over the signal intensity of the site-specific antibody. Phospho-ratios of an absolute fold change > 1.4 or < 0.4 were included in the analysis.

**Supplementary Table 2:**
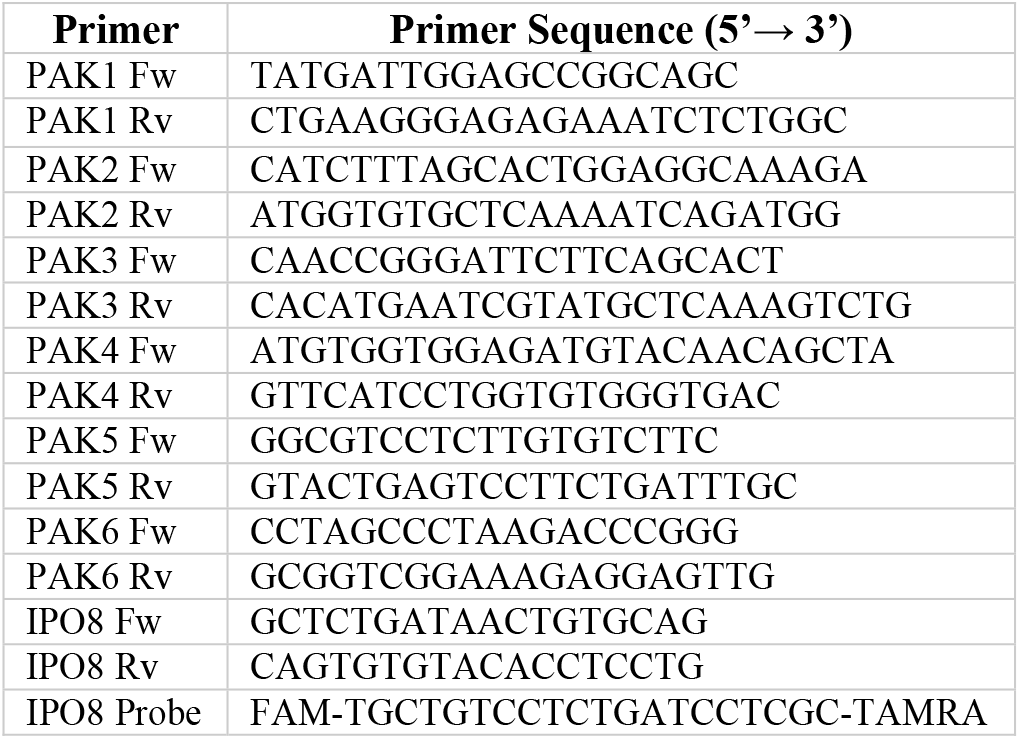
Primer sequences used for qPCR analysis of PAK isoform expression.

**Supplementary Figure 1:**
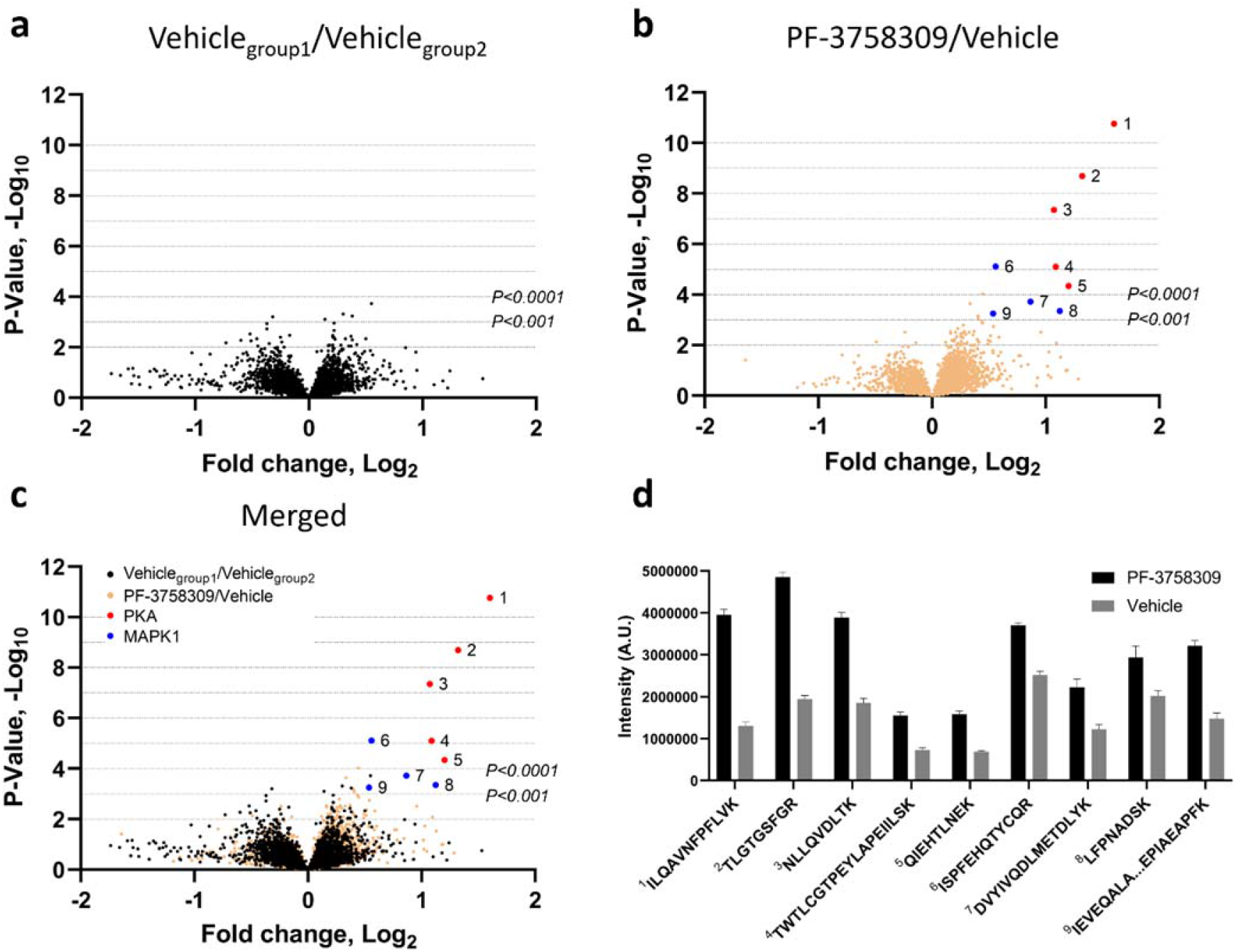
Drug/protein binding analysis of PF-3758309-treated peripheral blood mononuclear cells. Volcano plots for (a) two identical vehicle groups (decoy comparison), (b) PF-3758309 and vehicle groups (experimental comparison), and (c) the decoy and experimental comparisons merged. The x-axis indicates the relative fold change (log_2_-transformed expression values of peptides in the absence or presence of PF-3758309) and the y-axis indicates the -log10 of the p values. (**d**) Bar graph illustrating enrichment of PKA and MAPK1 peptides in PBMC lysates treated with PF-3758309. Bar graph data are shown as the mean ± standard error.

